# Effect of water temperature on the behavior of *Neptunea cumingii* and the histology, immune enzyme activity, and transcriptome of its gills and kidneys

**DOI:** 10.1101/2021.05.24.445381

**Authors:** Dandan Zhang, Xiaoyu Dong, Jianye Zhu, Jiacheng Yang, Ying Tian, Luo Wang, Junxia Mao, Xubo Wang, Yaqing Chang, Zhenlin Hao

## Abstract

The behavior of the marine snail *Neptunea cumingii* cultured at different temperatures (0, 4, 8, 12, 16 (control), 20, 24, and 28°C) and the histology, immune enzyme activity, and transcriptome of its gills and kidneys were studied using ecological and molecular methods. At 0°C, most of the snails shrank in size and did not eat during the first 6 h. At 28°C, snails also did not eat, death began to occur at 24 h. when the temperature was 8 °C and 12 °C, there was no significant difference in the mean feed intake of *N. cumingii* (P>0.05). And it was higher than the average conch intake under other temperature conditions. However, the histology of the gills and kidneys differed among test temperatures. At 0°C, the morphology of the gill pieces was difficult to judge. At 24°C, edema was present in the small gills, and at 28°C the small gills were severely deformed, the distance between the gill capillaries and the surrounding area was enlarged, and the gill tissue was severely damaged. Temperature increase or decrease from 16°C caused the columnar cells of the kidney to become shorter and more numerous. The total antioxidant capacity (T-AOC), catalase (CAT) and superoxide dismutase activities (SOD) of the gill and kidney differed significantly among the temperature conditions (p < 0.05). Transcriptome results showed that temperature change resulted in up-regulation of 2339 genes and down-regulation of 2058 genes in the gills, and in up-regulation of 2300 genes and down-regulation of 2060 genes in the kidneys of *N. cumingii*. Furthermore, the DEGs were subject to GO and KEGG enrichment analysis, and which showed that most of the DEGs in gill were involved in protein folding, defolding, translation, ribosome, and most of the DEGs in kidney were involved in DNA recombination, nuclear euchromatin, RNA-directed DNA polymerase activity. Finally, the results from this study show that *N. cumingii* prefers the temperature range was 8 to 16, temperature increase or decrease from that range, many aspects of *N. cumingii* biology could be significantly affected.

## 1. Introduction

*Neptunea cumingii* is a large warm water marine snail, and in China, it is mainly distributed in the subtidal zone of the Bohai and Yellow Sea areas (Zhou et al., 1995; Gao et al., 2015). *N. cumingii* has a spindle-shaped exterior shell with a bulged center and pointy ends as well as a horny operculum. It is carnivorous and feeds on benthic shellfish and rotten meat. *N. cumingii* is a popular and highly valued edible species due to its delicious and nutrient-rich meat, which contains a variety of amino acids, glycogen, protein, and essential trace elements that are good for human health (Cai, 2001).

To date, studies of *N. cumingii* have focused on its shell, distribution, and genetic diversity. For example, Zhao et al. (2004) studied the structural characteristics and the relationship between the structure and properties of the *N. cumingii* shell, and found that the shell of the conch Hemifusus tuba was composed of calcite and aragonite which were enchased in organic phase. Guo et al. (2008) surveyed the stock distribution of *N. cumingii* using bottom trawlers in Liaodong Bay, and found that the *N. cumingii* was mainly distributed in Laotieshan sea area of Lvshun, the relative biomass of *N. cumingii* and the bottom salinity had a positive correlation. Yu et al. (2019) introduced the reproductive biology of *N. cumingii* and the progress of artificial breeding technology. Azuma et al. (2009) studied the polymorphic microsatellite markers of *N. cumingii* in Hokkaido Japan, and isolated eight polymorphic microsatellite DNA loci. Their results suggested that the loci could be used as markers for population and kinship analyses in this species. Hao et al. (2020) studied the copulation, egg laying, embryonic development and changes in amino acids and fatty acids in *N.cumingii* during embryogenesis to understand the embryo development process and nutritional requirements in the early life phase. whose results showed that *N.cumingii* had direct development within the egg capsule and the development of embryos was classified into five stages. However, nothing is known about the effect of water temperature on the behavior, histology, immune enzyme activity, or transcriptome in the gill and kidney of *N. cumingii*. In this article, we studied the behavior, the histology, immune enzyme activity, and transcriptome of its gills and kidneys of *N*. *cumingii* cultured at different temperatures (0, 4, 8, 12, 16 (control), 20, 24, and 28°C) to obtain some useful information for the cultivation and protection strategies of the *N. cumingii*.

## 2. Materials and Methods

### 2.1 Experimental materials

*N. cumingii* were collected from the Lvshun Sea, Dalian City, Liaoning Province, China, and transported to the Key Laboratory of Mariculture and Stock Enhancement in North China’s Sea (Ministry of Agriculture, Dalian Ocean University, Dalian, P.R. China) in insulated containers. In the laboratory, samples were cultured in five 300L aquariums for two weeks before the experiments commenced. Each aquarium accommodated 95 *N. cumingii*, and the average individual wet weight was 85.3 ± 3.2 g. the temperature was 16 ± 1°C, and the salinity was 30 ± 1‰. Enough *Unionidae* (3 Kg) were thrown into each aquariums for *N. cumingii*’s food at 08:00 every day, 6 hours later, the residual bait and feces were clean up. The water was changed twice a day, half of the water was changed each time.

### 2.2 Experimental design

Before the experiment, all *N. cumingii* were deprived of food for two d. The eight temperature levels in the experiment were 0, 4, 8, 12, 16, 20, 24, and 28°C. 432 healthy and active *N. cumingii* were randomly selected and equally placed into 24 tanks (each temperature contained 3 tanks forming three replicates with 18 animals per tank). To achieve each test temperature, the temperature was increased or decreased from 16°C by 1°C/d. When the desired temperature was reached, that temperature was maintained for 72 h prior to the relevant experiments.

#### 2.2.1 Behavior

To assess behavior, 80 active *N. cumingii* (10 per temperature) were placed in eight observable containers for 2 h, and then 1 kg of live *Unionidae* were added to the container. The feeding behavior of *N. cumingii* was observed for 6 h. Feeding rate was calculated as the followings:

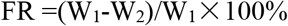

where W_1_ and W_2_ are the initial and final wet weights of *Unionidae.*

#### 2.2.2 Histology

Three individuals per replicate were selected for histological and enzyme activity assays. Fresh gill and kidney tissues were removed from *N. cumingii* and fixed in Bouin’s fixative solution for more than 24 h. Fixed specimens were preserved in 70% alcohol, then individually embedded in paraffin blocks and sectioned in serial sagittal sections (4 μm) using a rotary microtome (Leica, Germany). Tissue sections were permanently mounted, and the slides were stained with hematoxylin-eosin (H&E) for general histological observations. The sections were examined under Leica DM500 compound microscope. Microphotographs were taken with Leica MC170 HD camera.

#### 2.2.3 Enzyme activity

The activities of catalase (CAT) and superoxide dismutase (SOD) and total antioxidant capacity (T-AOC) levels in the gill and kidney of *N. cumingii* were measured using assay kits purchased following the manufacturer’s protocols. Enzyme determination was carried out at 25 ± 1°C in an air-conditioned room. T-AOC was measured based on the generation of the Fe^2+^-o-phenanthroline complex, as the overall reducing agents in the sample supernatant reduced Fe^3+^ to Fe^2+^, which reacted with the substrate o-phenanthroline. Stable color of the Fe^2+^ o-phenanthroline complex was measured at 520 nm at 37°C. One unit of T-AOC was defined as the amount necessary to increase the absorbance by 0.01 at 37°C (U mg^−1^ protein). The decomposition reaction of H_2_O_2_ by CAT can be terminated immediately by adding ammonium molybdate, which reacts with the remaining H_2_O_2_ to form a faint yellow complex. CAT activity was detected by measuring the decrease in absorbance resulting from H_2_O_2_ decomposition at 405 nm. One unit of CAT activity was defined as the amount of enzyme that decomposes 1 μmol of H_2_O_2_ (U/mg protein). SOD activity was determined at 550 nm using the xanthine and xanthine oxidase systems. One unit of SOD activity was defined as the amount of enzyme required to cause 50% inhibition of the xanthine and xanthine oxidase reaction in 1 ml of enzyme extract of 1 mg protein (U/mg protein).

#### 2.2.4 Transcriptome of gill and kidney

According to the previous results of behavior, histology and enzyme activity, we selected two significantly different treatment groups (16 °C and 28 °C)for transcriptome assay. The gill and kidney samples of *N.cumingii* at different test temperature (3 individuals per replicate) were collected and frozen in liquid nitrogen and stored in a refrigerator at −80°C for later use. Total RNA from gills and kidneys tissue of *N.cumingii* was extracted using the Trizol kit (Invitrogen, USA). Genomic DNA was treated with DNase I (TaKaRa, Dalian, P R China), after with 0.7% agarose gel electrophoresis was used to assess RNA integrity, with an Agilent 2100 Bio-analyzer (Agilent Technologies, USA) and ND-2000 (NanoDrop Technologies) platforms used to confirm RNA purity and quantity. Equal amounts of high-quality RNA (OD260/OD280 = 2.04−2.07, OD260/OD230 ≥ 1.65−2.18, RIN ≥ 8.4) from the gills and kidneys of snails were then pooled to construct a se-quencing library. mRNA was collected using oligo(dT) beads, followed by fragmentation buffer-mediated fragmentation. Random hexamer primers were used to generate double-stranded cDNA. After end repair, “A” base addition, and adapter connection, 200−300 bp cDNA frag-ments were selected and purified via agarose gel electrophoresis, am-plified by 15 PCR cycles, the library sequencing was conducted. Functional annotation and classification were conducted Blast2 GO and WEGO respectively. DEGs were compared with the terms in GO database to get the list and number of transcriptions belonging to the GO function. Then hypergeometric tests were used to identify GO entries in which DEGs were significantly enriched compared to the entire set of transcripts. KEGG is another leading public database about the pathways used for understanding high-level functions and utilities of the biological system. When DEGs were compared with the pathways in KEGG database, hypergeometric tests were used to identify the KEGG Pathways in which DEGs were significantly enriched. These GO terms and pathways that had Q-values ≤ 0.05 were deemed significantly enriched. Analysis based on GO terms and pathway is helpful to further understand the biological functions of transcripts. R was utilized for all the expression data statistics and visualization. Systematically analyze the data and compare the gills and kidneys of the snails.

### 2.3 Statistical analysis

SPSS 22.0 and OriginPro 2017 software were used to analyze the differences in snail feeding rate caused by temperature changes over a 6 h period.

STATISTICA6.0 software was used to analyze the enzyme activity data. The results were expressed as the mean ± standard error. Significant differences between groups were analyzed by one-way analysis of variance (ANOVA) and Tukey’s honestly significant different test where appropriate. p < 0.05 was considered statistically significant (95% confidence interval).

Statistical analysis of transcriptome data was performed using Excel 2016 and SPSS 19.0. All data were presented as the mean ± standard deviation. Significant differences were analyzed via one-way ANOVA, and p < 0.05 was considered statistically significant. Pearson correlation analyses were used to assess the strength of the relationship between RNA-Seq data and RT-qPCR measurements.

## 3. Results

### 3.1 Behavior

Behavior of *N. cumingii* differed among the different temperature conditions (Figure 1, Table 1). At 0°C, most of the *N. cumingii* shrank in size and did not eat during the first 6 h. At 28°C, the snails also did not feed. However, death began to occur at 24 h, and all snails died within 72 in the 28°C group. There was no significant difference in the behavior of the *N. cumingii* when the temperature was 4–24°C, but food intake did differ significantly. The food intake at 16°C was significantly higher than at the other temperatures, followed by 8 and 12°C, which did not differ significantly from each other.

**Figure 1.**
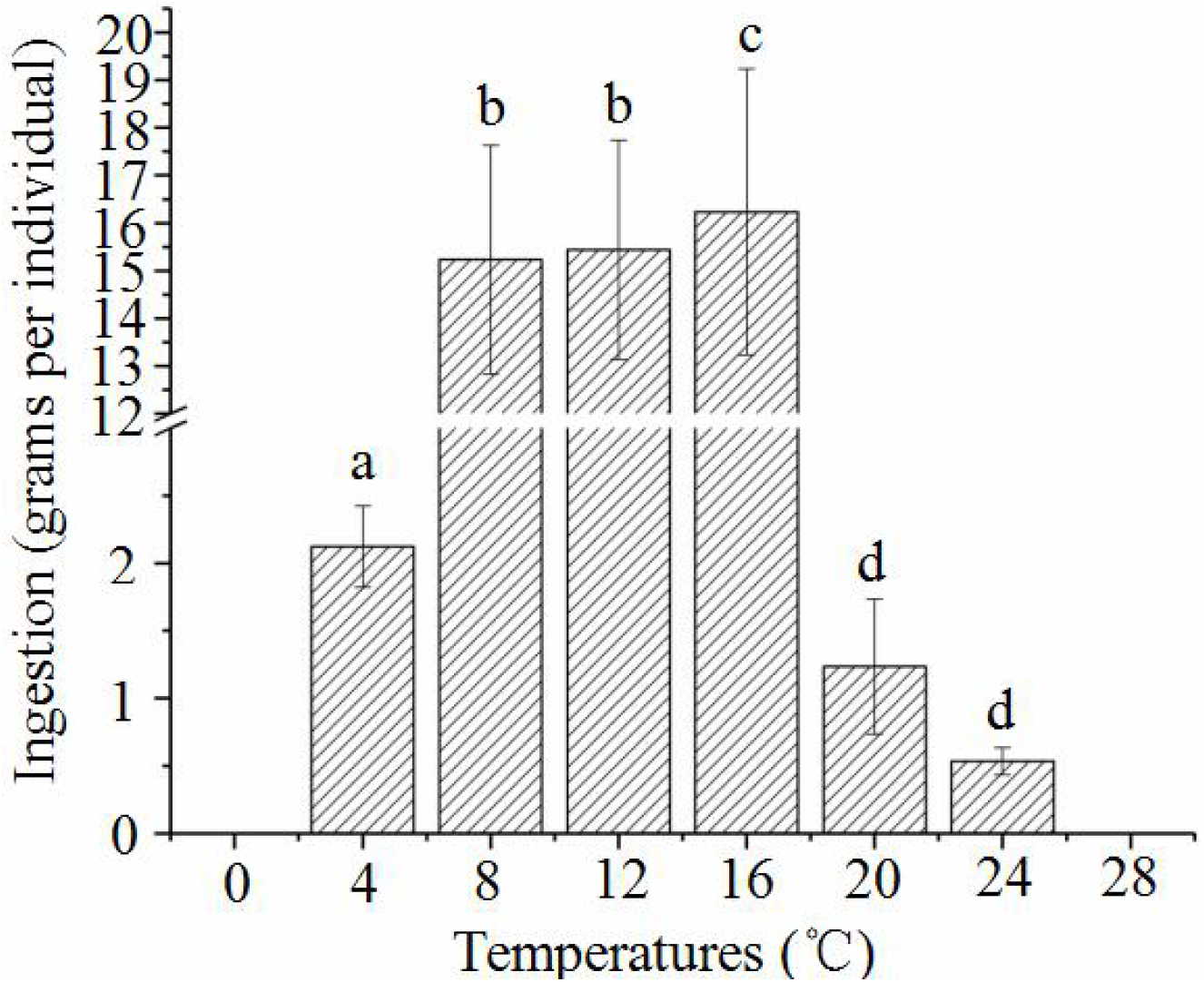
*N. cumingii* feeding behavior when cultured at different temperatures for 6 h. Different lowercase letters indicate significant differences in feeding rates of *Neptunea cumingii* at different temperatures (p < 0.05).

**Table 1.**
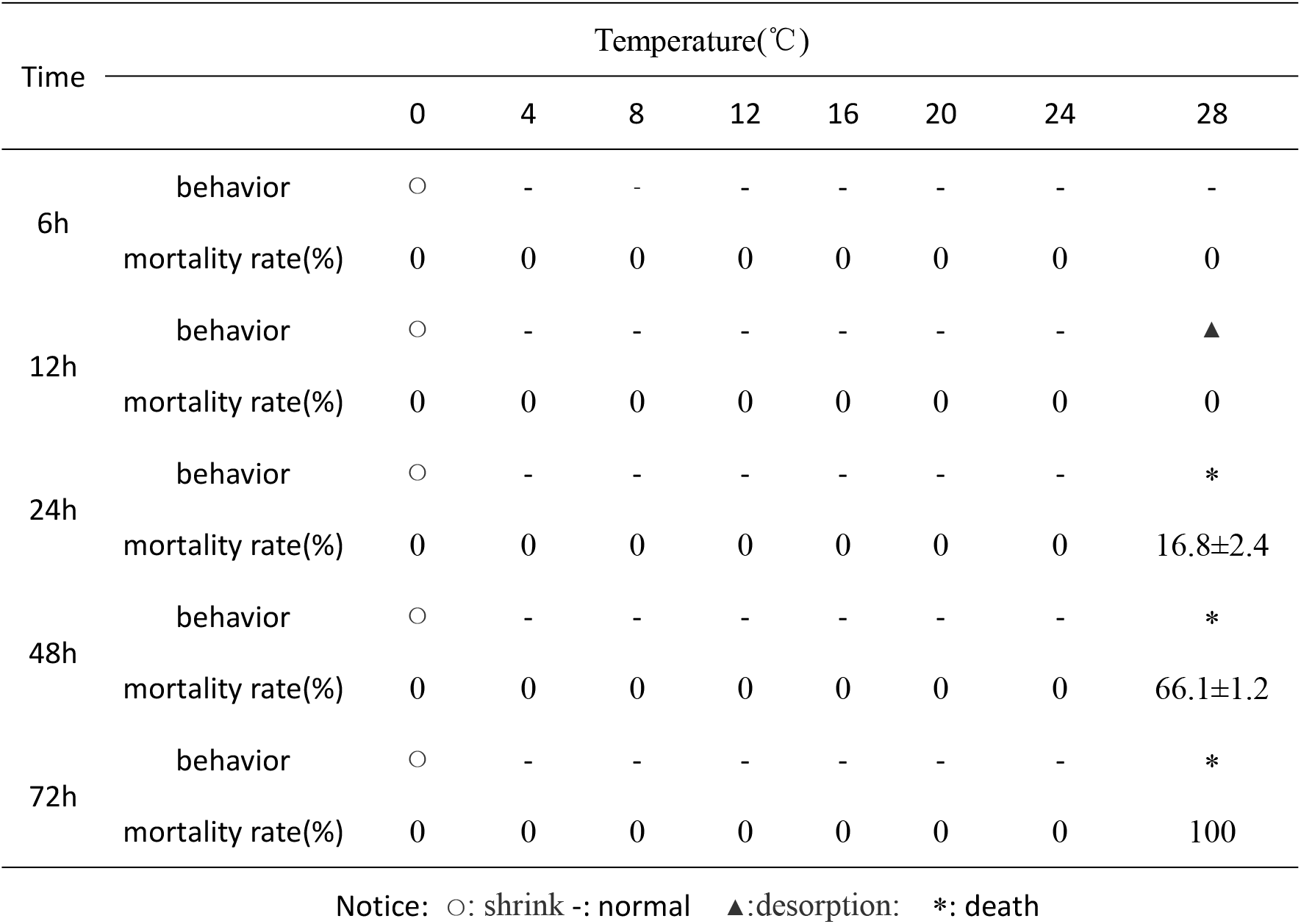
Effects of different temperatures on the behavior and mortality of *Neptunea cumingii*

**Table 2.** Annotation information statistics for the gill and kidney transcriptome of *Neptunea cumingii*

|  | Nr note | Swissprot | KEGG | KO notes | eggNOG | GO notes | Pfam |
| --- | --- | --- | --- | --- | --- | --- | --- |
|  |  | notes | notes |  | notes |  | notes |
| Number of genes | 20763 | 13106 | 9070 | 11068 | 15224 | 12015 | 12293 |
| Proportion (%) | 25.50 | 16.10 | 11.14 | 13.59 | 18.70 | 14.76 | 15.10 |

Shrink: The rhynchodaenm is in the body without extending, and the foot extension area is very small. Normal: The foot extension area is larger than that in shrink condition, and the rhynchodaenm is out of the body. Desorption: The absorption force of the foot becomes so weak that they often failed to hold onto the wall of the pool. Death: The foot and operculum turn out of the shell without shrinking, even when they are touched with fingers.

### 3.2 Histology of gills and kidneys in *N. cumingii*

At the control temperature of 16°C (Figure 2E), a blood vessel was visible in the center of the gill filaments, and small gills were present on both sides of the gill filaments to allow gas exchange. Mitochondria-rich cellswere distributed at the base of the gill lamella. At 8°C, the gill filaments were swollen and numerous blood cells flowed out of them. At 4°C, the epidermis of the gill lamella exhibited shedding, and the distance between adjacent gill lamellas was enlarged. At 0°C, the morphology of the gill tissue was difficult to judge because the low temperature had destroyed much of the tissue. At 24°C, edema was visible in the small gills, and at 28°C the small gills were severely deformed, the distance between the gill capillaries and the surrounding area was enlarged, and the gill tissue was severely damaged. (Figure 2)

**Figure 2.**
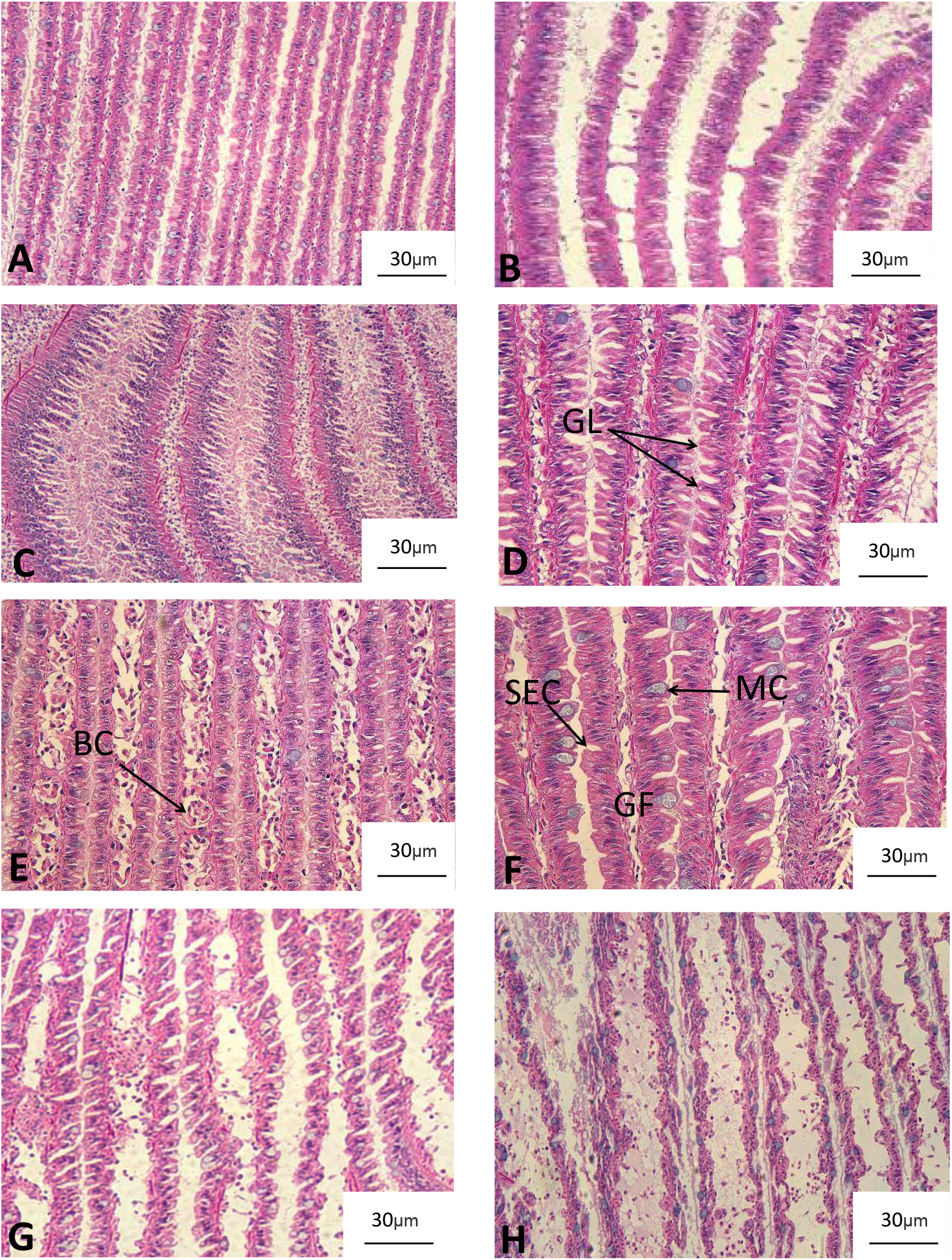
Histology of the gills of ***Neptunea cumingii*** cultured at different temperatures. A: 0°C, B: 4°C, C: 8°C, D: 12°C, E: 16°C, F: 20°C, G: 24°C, H: 28°C. GL: gill lamellae, BC: blood corpuscle, SEC: squamous epithelial cells, MC: mucous cells, GF: gill filament

The kidneys of *N. cumingii* contained only tubules and collecting ducts. The kidney columnar cells in the 16°C group were longer than those in the other groups; at both higher and lower temperatures, the columnar cells were shorter and more numerous. (Figure 3)

**Figure 3.**
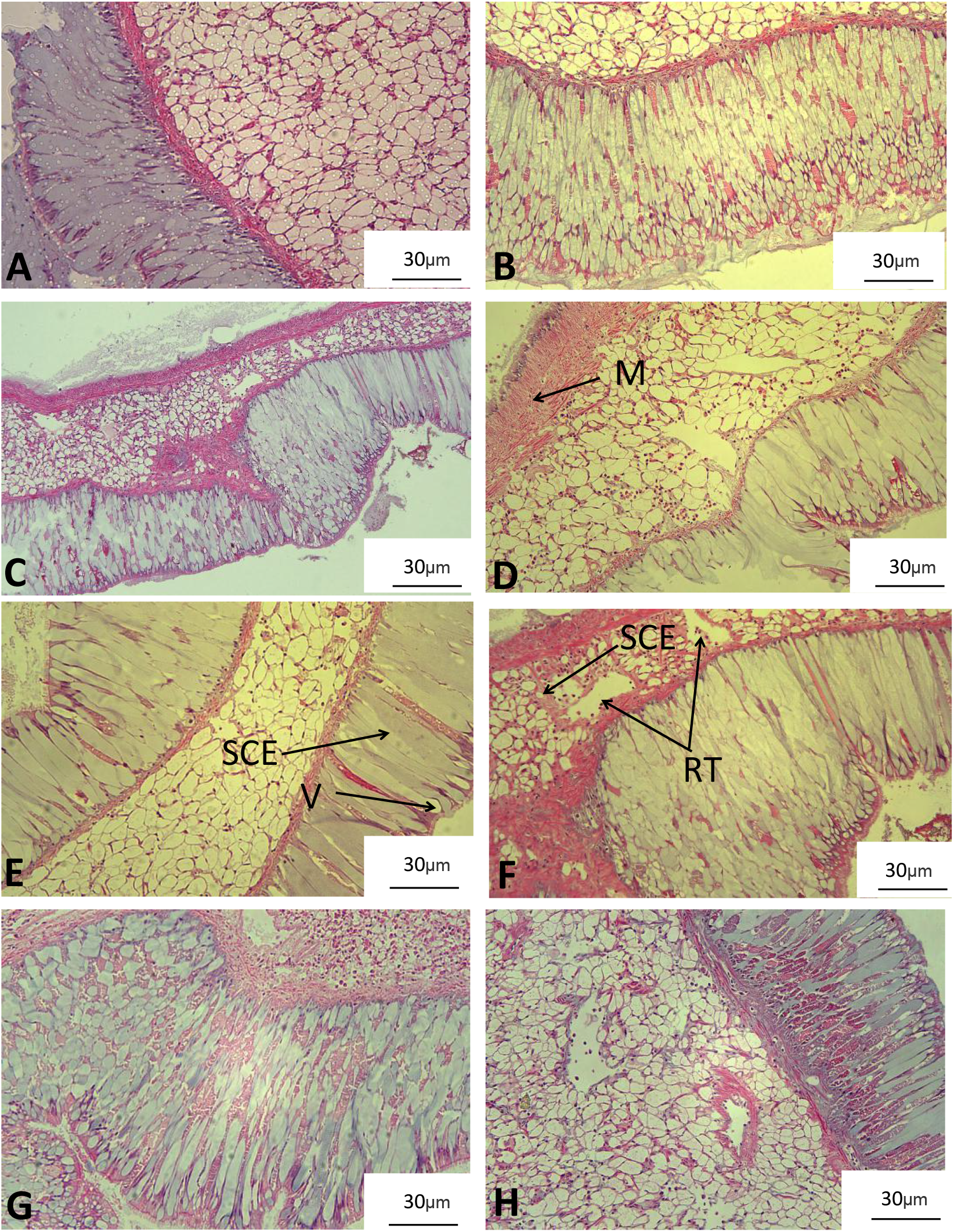
Histology of the kidney of *Neptunea cumingii* cultured at different temperatures. A: 0°C, B: 4°C, C: 8°C, D: 12°C, E: 16°C, F: 20°C, G: 24°C, H: 28°C. CT: collecting tube, RT: renal tubule, M: muscle, V: villi, SCE: simple columnar epithelium

### 3.3 T-AOC and activities of CAT and SOD

Figure 4 shows the T-AOC of the gills and kidneys of *N. cumingii* cultured under different temperature conditions. At temperatures ranging from 4 to 24°C, the T-AOC of the gill was maintained between 0.7 and 1.0 U/mg prot, and values did not differ significantly between the 16 and 20°C groups (p > 0.05). At 0 and 28°C, the T-AOC of the gills was low. There was no significant difference in T-AOC of the kidney among the 12, 16, and 20°C groups (p > 0.05), but the values gradually decreased as temperature increased to 24 and 28°C. Additionally, T-AOC of the kidney increased from 0 to 4°C and was highest at 8°C.

**Figure 4.**
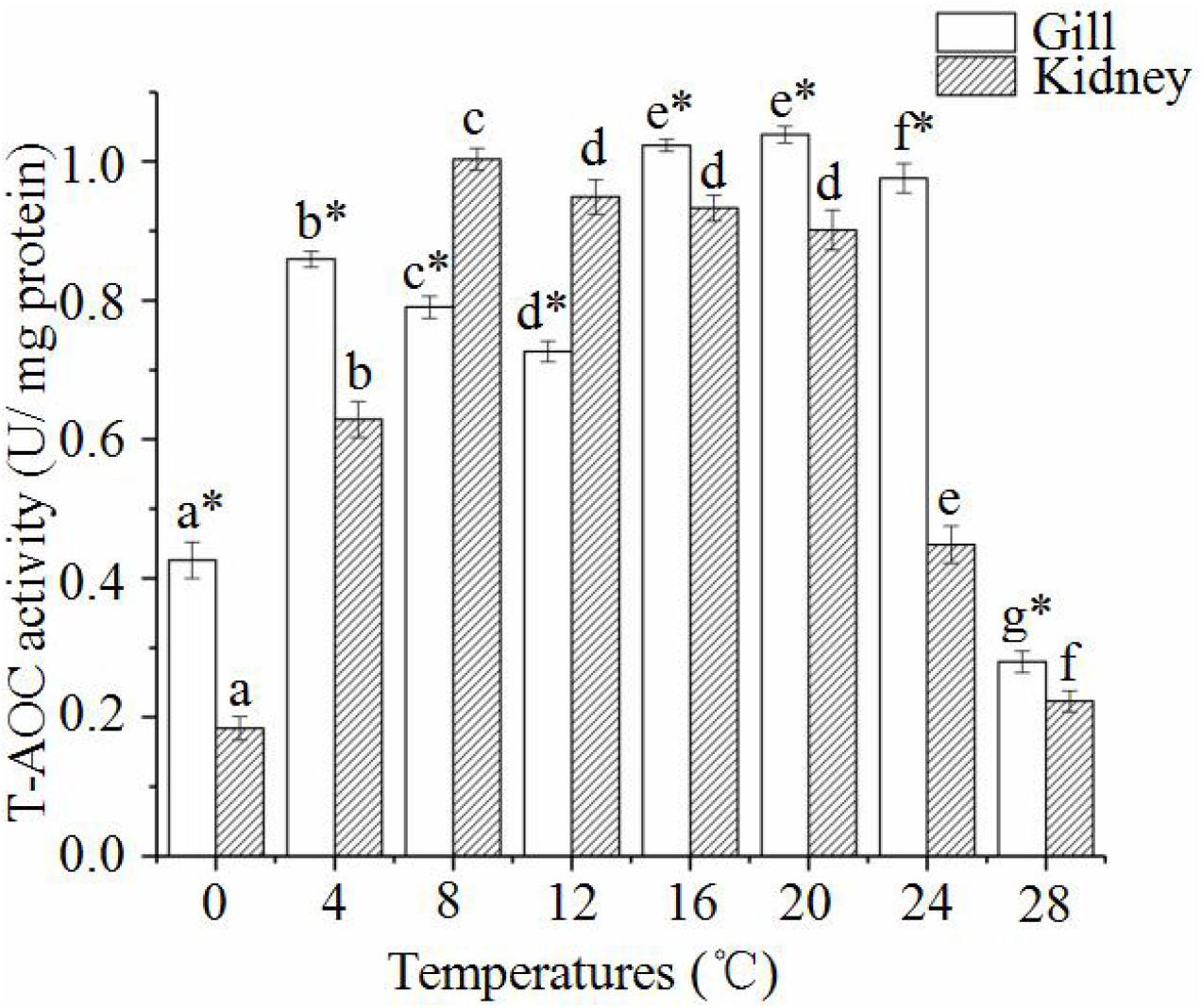
T-AOC of the gills and kidneys of *Neptunea cumingii* cultured at different temperatures. Letters indicate significant differences of T-AOC activities among the treatments (P<0.05). * indicate significant differences on T-AOC activities between the gill and kidney (P<0.05).

The CAT activities of the gills and kidneys of *N. cumingii* cultured at different temperatures are shown in Figure 5. At 0°C, the CAT activity of the gill was 0.59 U/mg prot, but it was lower (0.30 U/mg prot) in the 4, 8, and 12°C groups and then higher (0.65 U/mg prot) in the 16, 20, and 24°C groups. The activity was lowest (0.23U/mg prot) in the 28°C group. CAT activity of the gills was much lower than that of the kidney at all temperatures. The activity increased and decreased several times with increasing culture temperature, but the values did not differ significantly between the 0 and 4°C groups, between the 8 and 12°C, or between the 16 and 20°C groups (p > 0.05). The lowest activity occurred in the 28°C group.

**Figure 5.**
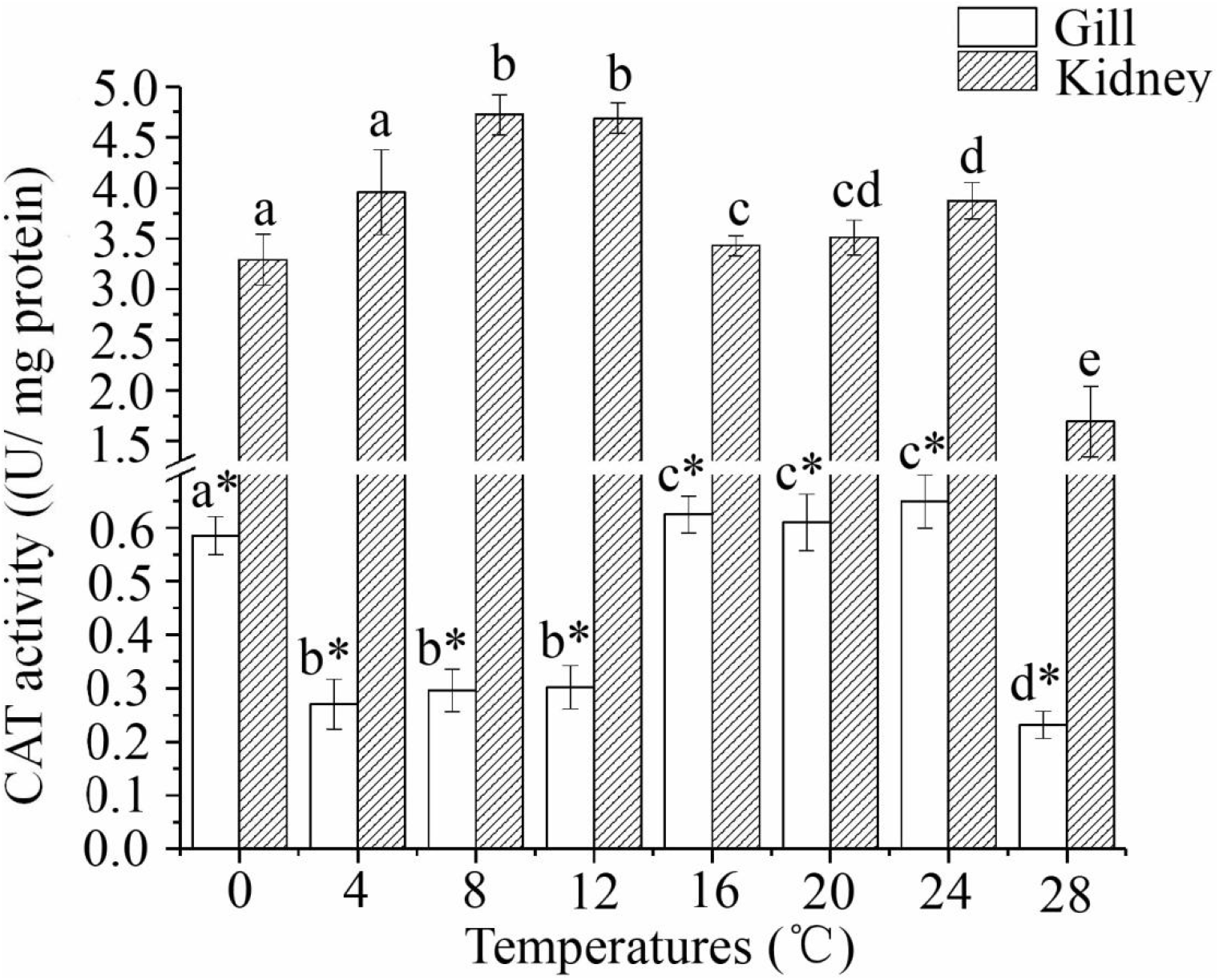
CAT activity of the gills and kidneys of *Neptunea cumingii* cultured at different temperatures. Letters indicate significant differences of CAT activities among the treatments (P<0.05). * indicate significant differences on CAT activities between the gill and kidney (P<0.05).

Figure 6 shows the SOD activities of the gills and kidneys of *N. cumingii* cultured at different temperatures. As the temperature increased from 0 to 28°C, the activity of SOD in the kidney exhibited a W-shaped pattern. The SOD activity of the gills was maintained above 150U/mg prot at all temperatures. In the kidney, the highest SOD activity (120 U/mg prot) occurred in the 0°C group. SOD activity differed significantly among all temperature groups (p < 0.05) except for the 4, 8, and 20°C, which did not differ from each other (p > 0.05).

**Figure 6.**
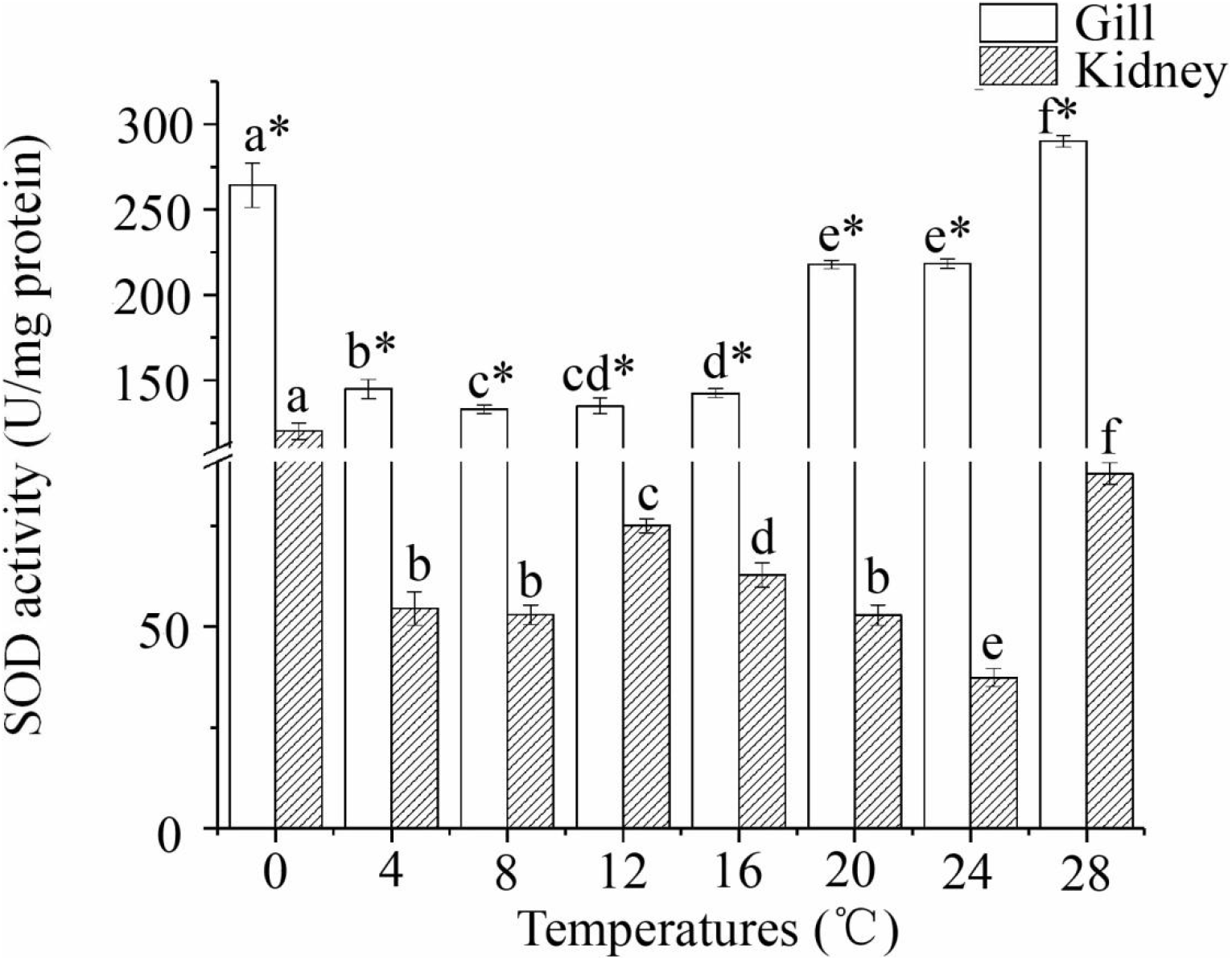
SOD activity of the gills and kidneys of *Neptunea cumingii* cultured at under different temperatures. Letters indicate significant differences of CAT activities among the treatments (P<0.05). * indicate significant differences on CAT activities between the gill and kidney (P<0.05).

### 3.4 Transcriptome analysis

Transcriptome data showed that the original data Q30 of each sample were distributed in 91.64–92.25%, the effective data volume was distributed in 6.12–6.89 G, and the average GC content was 45.04%. In total, 81,429 unigenes were spliced together, with a total length of 67,628,134 base pairs and an average length of 830 base pairs. Table 1 shows the unigene database annotation results. The comparison rate of reads to unigenes was 83.53–84.33%. The numbers of differentially expressed genes detected were 4397 for gills and 4360 for kidneys. In total, 52,399 simple sequence repeats (SSRs), 29,251 unigenes containing SSRs, 12,391 unigenes containing more than 1 SSR, and 11,640 composite SSRs were predicted. Additionally, 40,930 coding sequences were predicted (20,776 were predicted by the database comparison method and 20,154 were predicted by ESTScan).

Analysis showed that the assembled unigenes participate in 42 functional pathways (Figure 7) in the following six categories: cellular processes, environmental information processing, genetic information processing, metabolism, human diseases, and organic systems. The number of genes in the different metabolic pathways varied greatly.

**Figure 7.**
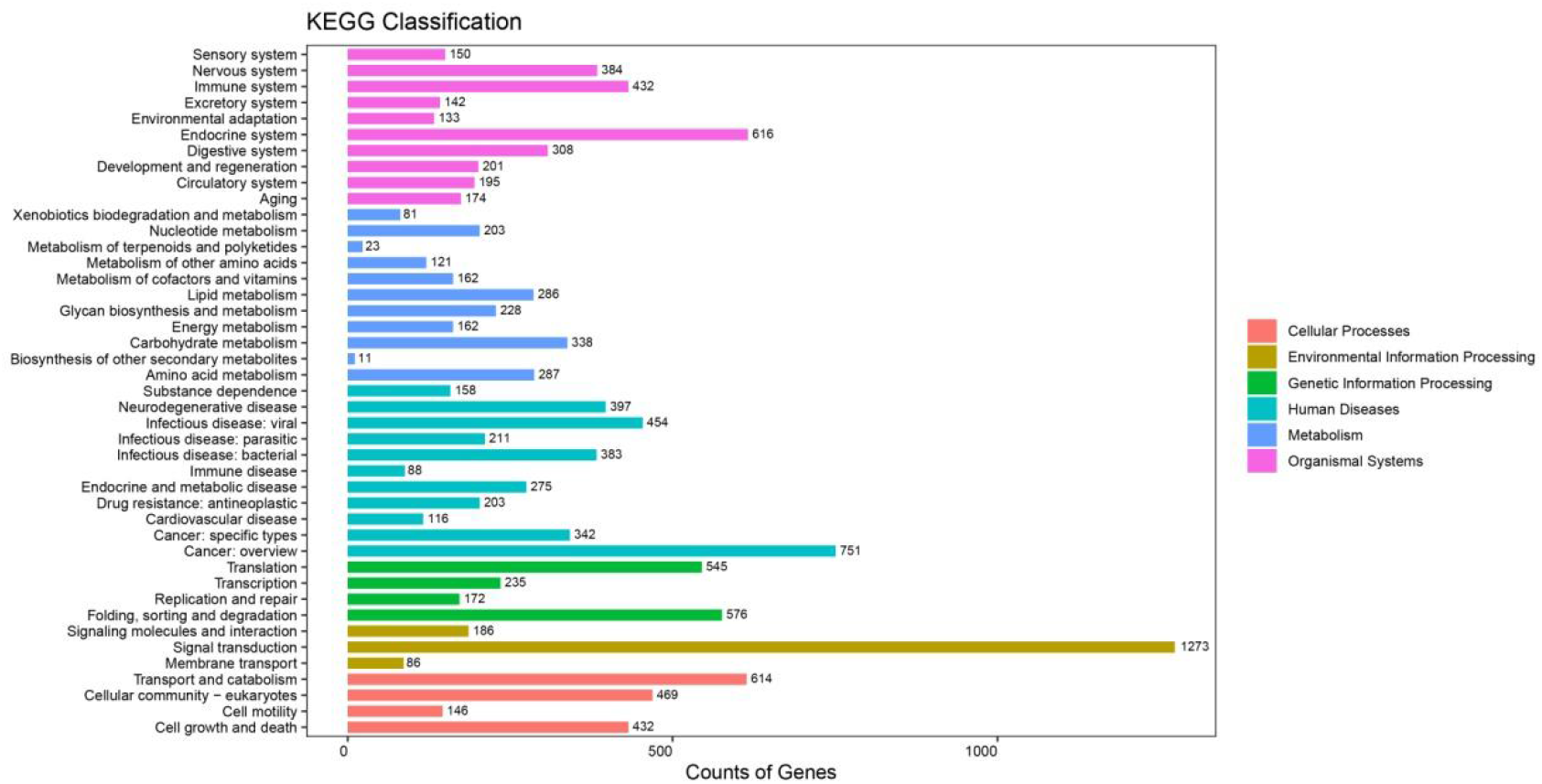
KEGG annotation statistics chart for the gill and kidney. The horizontal axis represents the number of genes, the vertical axis represents the name of the Level2 pathway, and the number on the right side of the column represents the number of genes annotated to the Level2 pathway.

KOG annotation of the unigenes was performed to classify them into KOG groups (Figure 8). The letters on the horizontal axis indicate the names of the 25 KOG groups, and the ordinate shows the proportion of the number of genes annotated to this group to the total annotated genes. Gene function prediction was annotated the most times, followed by signal transduction mechanism. Post-translational modification, protein conversion, and molecular chaperones also were annotated numerous times, whereas cell activity was annotated least, which meets the experimental conditions.

**Figure 8.**
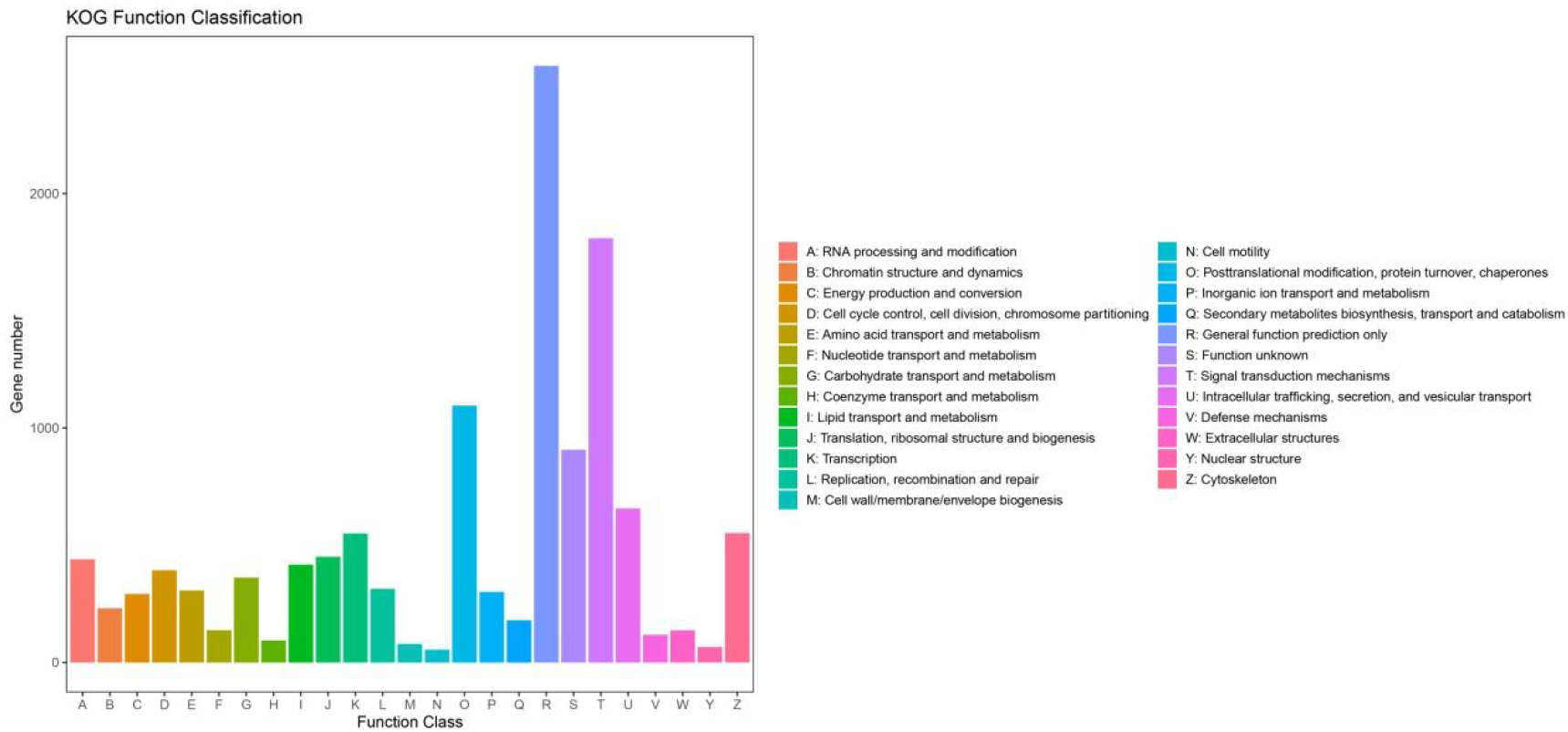
KOG function classification diagram for the gill and kidney. The horizontal axis represents the functional classification of KOG, and the vertical axis represents the number of genes.

After initial GO annotation of the unigenes, the successfully annotated genes were classified according to the next level of the three major categories of GO, which were biological process, cell composition, molecular function (Figure 9). The abscissa shows the GO term of the next level of each of the three major categories of GO and the ordinate shows the number of genes annotated to the term. There were relatively more annotated genes for cellular processes, metabolic processes, and single biological processes in the biological process category. For the cell composition category, cells, cell parts, and organelles had the most annotated genes, and for the molecular function classification, the most annotated genes were in the connection and catalytic activity categories.

**Figure 9.**
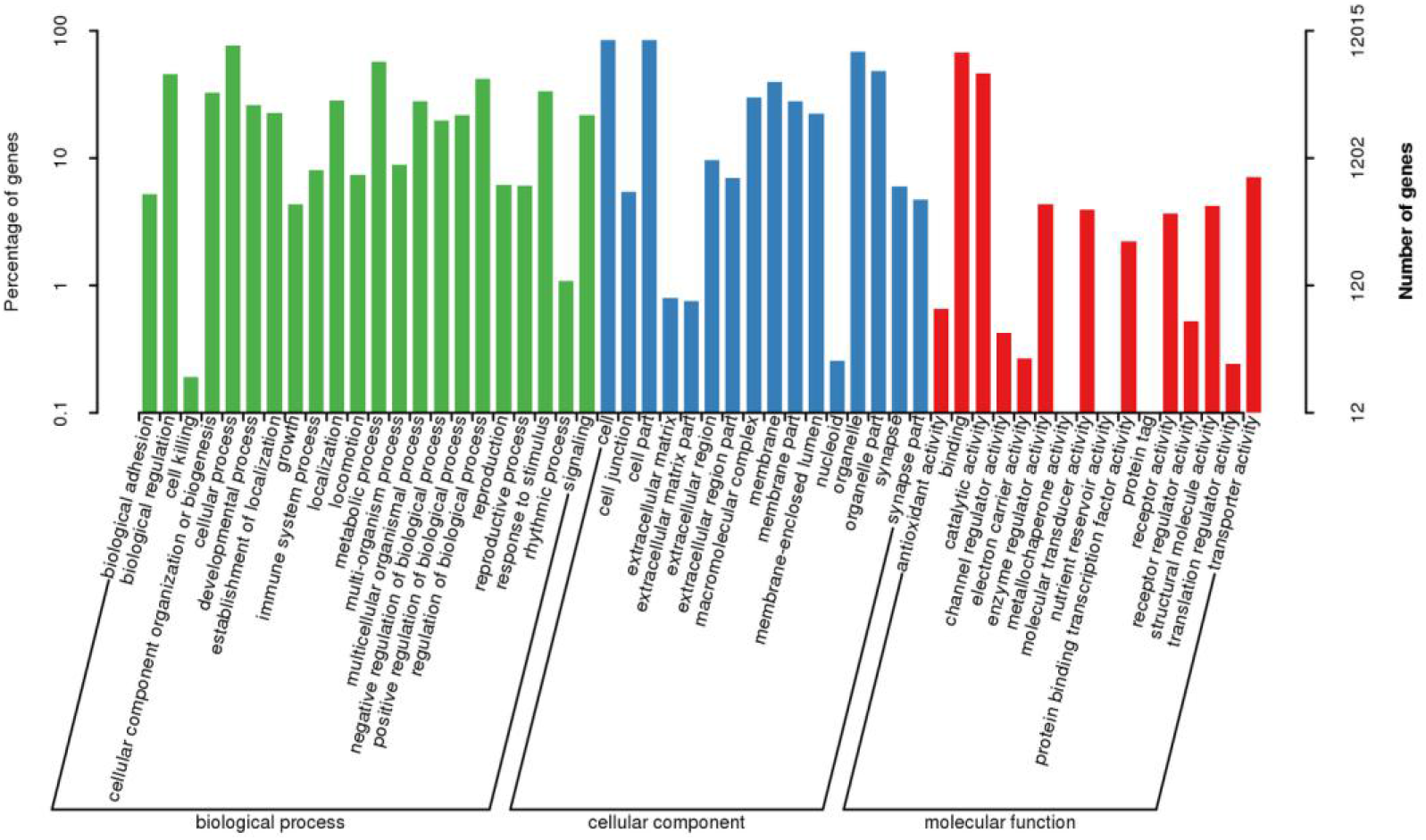
GO function classification diagram for the gill and kidney. The horizontal axis represents the GO function classification, the left vertical axis represents the proportion of genes annotated to this category, and the right vertical axis represents the number of genes annotated to this category.

As shown in Figures 10 to 12, 4397 genes in the gills of *N. cumingii* showed differential expression (2339 up-regulated and 2058 down-regulated), and 4360 genes in the kidney showed expression differences (2300 up-regulated and 2060 down-regulated). In GO enrichment, DEGs of gill significantly enriched in protein folding, translation, ribosome, unfolded protein binding and structural constituent of ribosome terms. And DEGs of kidney significantly enriched in DNA recombination, nuclear euchromatin, RNA - directed DNA polymerase activity and aryl sulfotransferase activity terms. |Log2fc | > 1 and FDR < 0.05 were used as the selection standards for differential expression in KEGG enrichment analysis. We selected the first 20 pathways with significant enrichment and found the DEGs of gill significantly enriched in Ribosome, Protein processing in endoplasmic reticulum, Apoptosis, NOD-like receptor signaling pathway and TNF signaling pathway. The Protein processing in endoplasmic reticulum involved the most DEGs (24 DEGs) in the gill, which also involved Hsp 70 and Hsp 90. The DEGs of kidney significantly enriched in NF-kappa B signaling pathway, Longevity regulating pathway – multiple species, Apoptosis – multiple species, NOD–like receptor signaling pathway and TNF signaling pathway. The most DEGs involved in TNF signaling pathway(12 DEGs).

**Figure 10.**
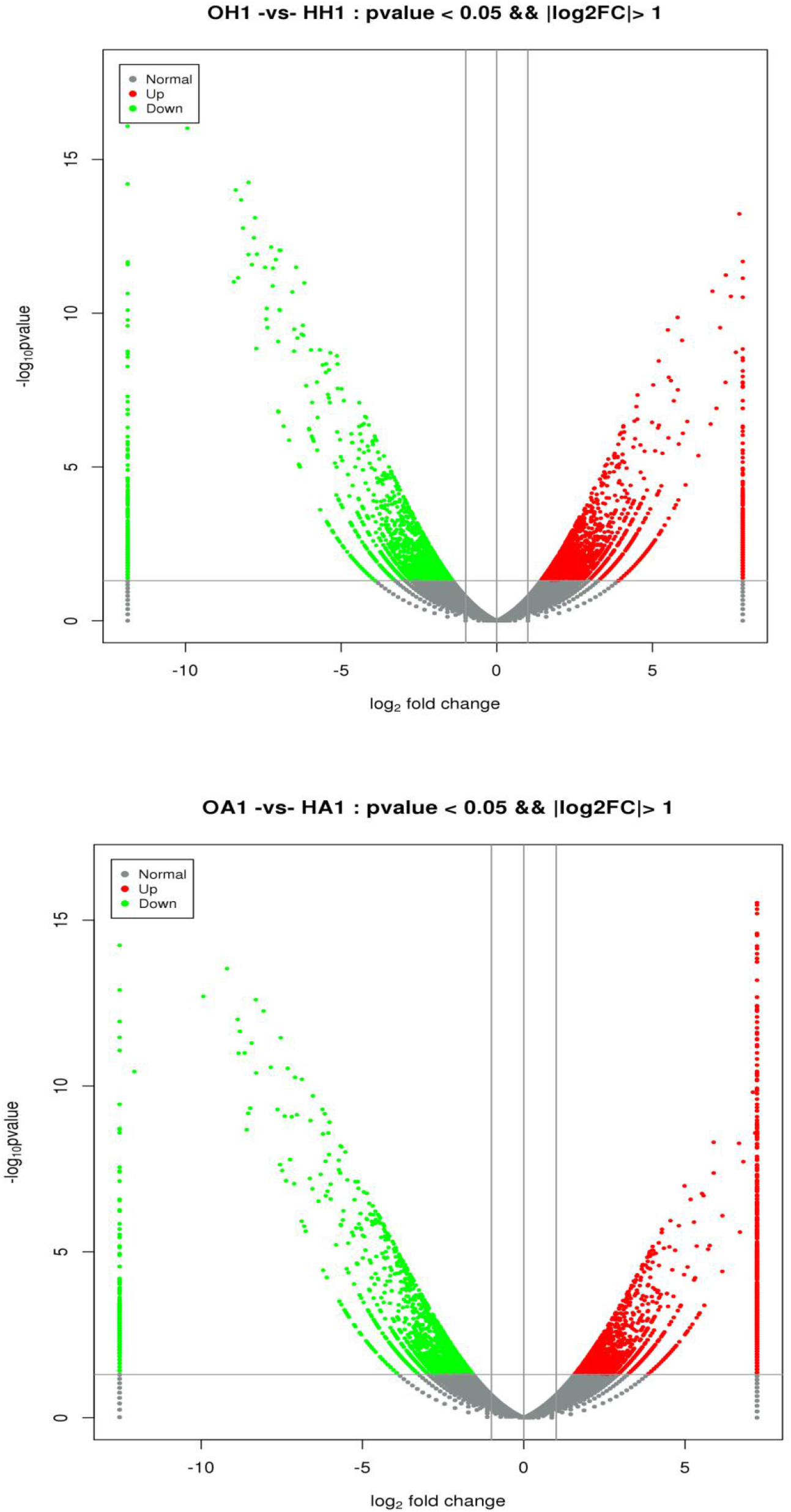
Differential gene comparison volcano map for the kidney (above) and gills (under). Gray is non-difference Unigene, red is up-regulated significant difference Unigene, and green is down-regulated significant difference Unigene; The X-axis is log2 FoldChange, and the Y-axis is log10Pvalue.

**Figure 11.**
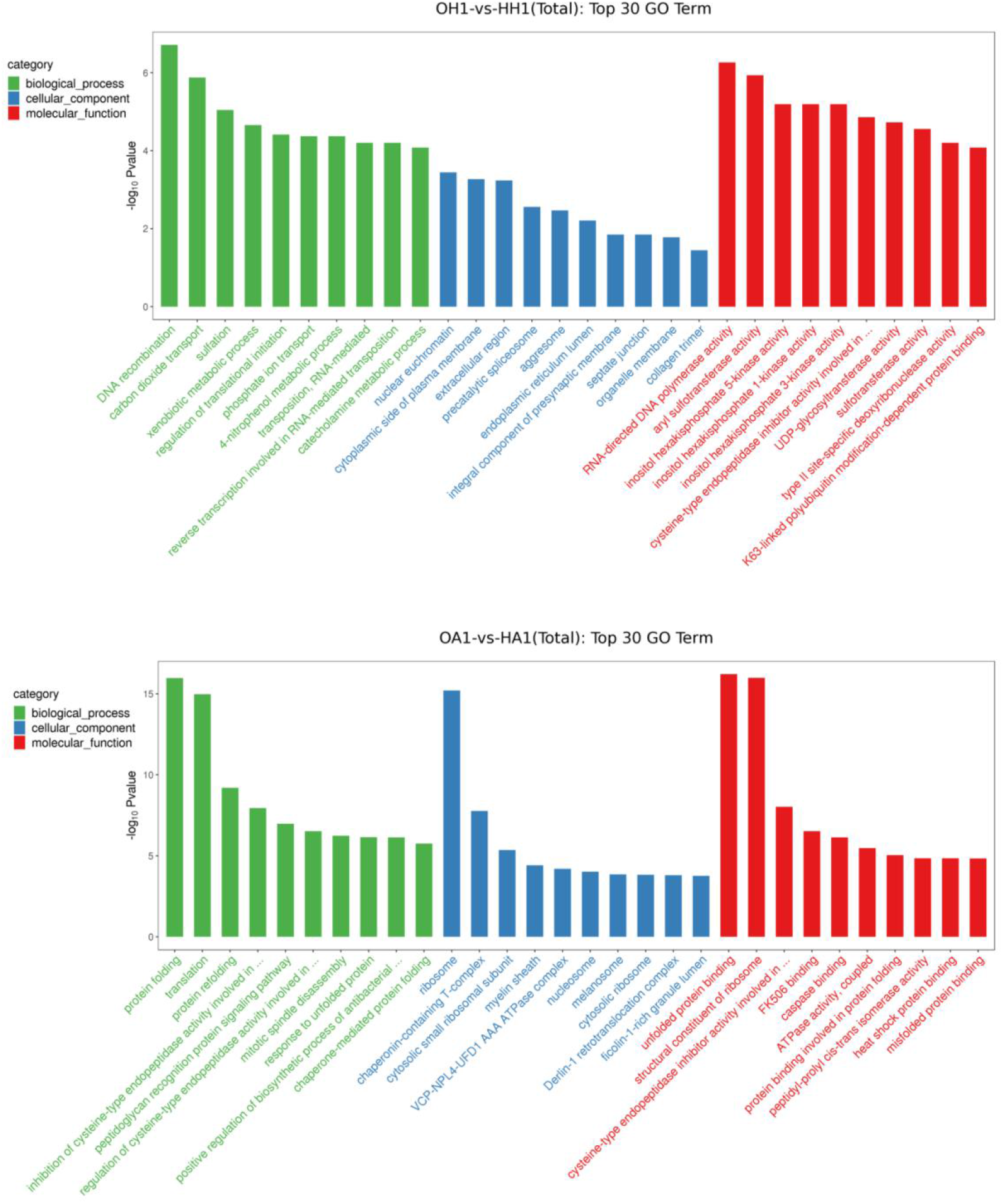
GO enrichment map for the kidney (above) and gills (under) The X-axis is the name of the GO item, and The Y-axis is −log10Pvalue.

**Figure 12.**
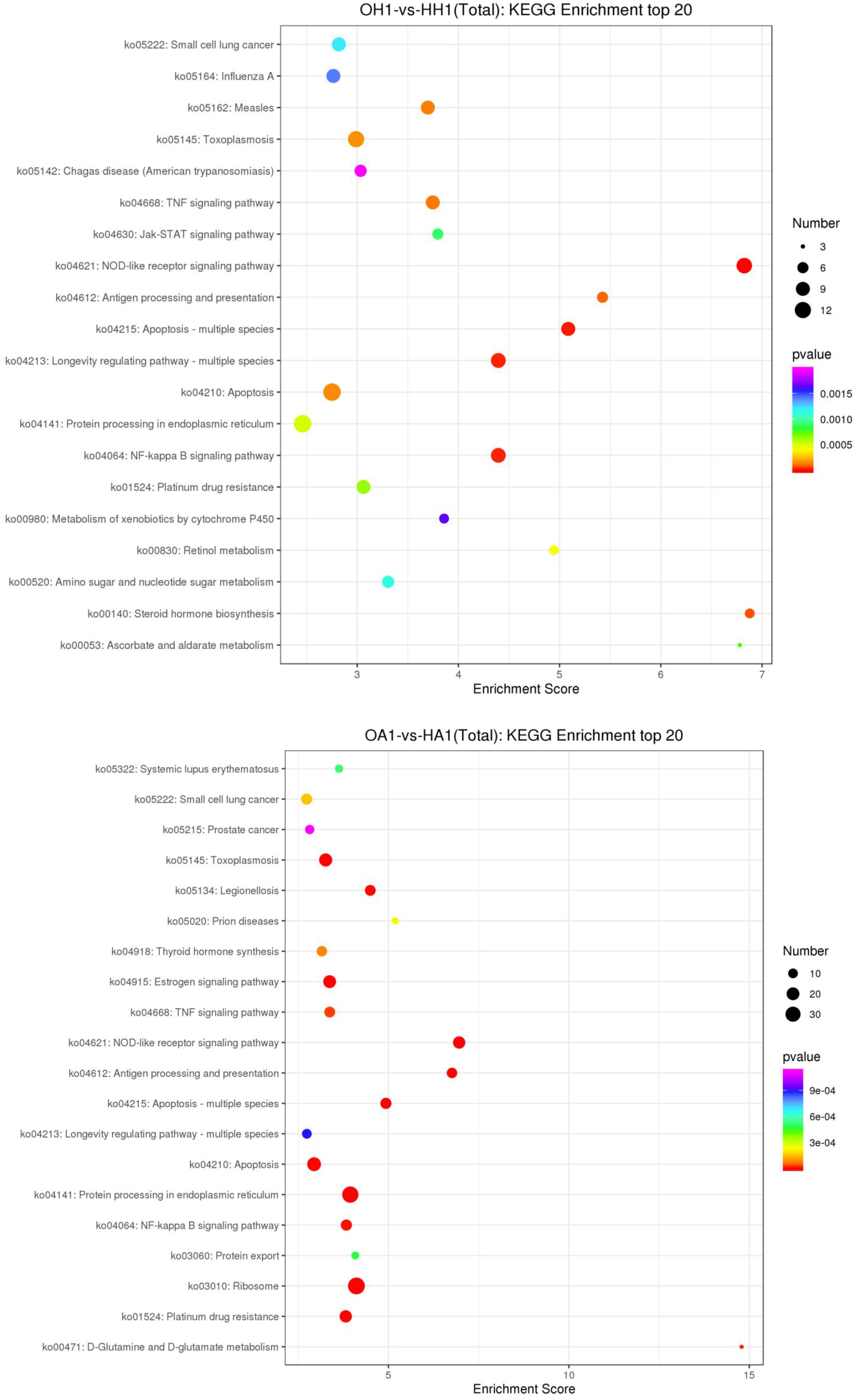
KEGG enrichment map for the kidney (above) and gills (under) The X - axis is the enrichment score. The larger the bubble, the more the number of different Unigenes, and the color of the bubble changes from purple-blue-green-red. The smaller the enrichment pvalue, the greater the degree of significance.

## 4. Discussion

### 4.1 Behavior

Water temperature has significant effects on movement, feeding, reproduction, and behavior of aquatic animals. For example, Keen et al. (1994) showed that the swimming speed of *oncorhynchus mykiss* increased proportionally with increasing temperature but then decreased when the temperature exceeded the optimum value. Chen (2004) found that the food intake of *Cyprinus carpio* first increased and then decreased with increasing culture temperature. In another study, Armstrong et al. (2013) found that *Oncorhynchus keta* spawning grounds were situated only in areas with suitable water temperature. In this study, we also found that temperature had a significant impact on the behavior of *N. cumingii*. When the temperature was very low (0°C), the snails contracted, closed their shells, and did not eat. As temperature increased, food intake first increased, peaked at 16°C, and then decreased. The highest average food intake was 16 gram per individual, which was consistent with the normal food intake of *N. cumingii*. (Fujinaga et al., 1999; Miranda et al., 2008). Other studies have shown that an initial increase followed by a decrease in food intake as temperature increases is a common phenomenon in aquatic animals (Sun et al., 1982; Xie et al., 1992). Li (2006) reported that digestive enzyme reactions in the black owl body accelerated with the increase in temperature within a certain range, but after a certain value the speed of the reactions slowed down. We propose that the weakened feeding ability of *N. cumingii* that occurred at 28°C may have been due to the decreased concentration of dissolved oxygen in the water at higher temperature. This would result in more energy being used to maintain resting metabolism and less energy available for food digestion and thus decreased food consumption by *N. cumingii*. Similar results also occurred in *Mizuhopecten yessoensis* (Ben, 2013) and *Haliotis discus hannai* (Jiang, 2017). When the temperature was 28°C, the snails had poor adsorption ability and gradually closed their shells within 12 h. Death began to occur at 24 h, and all snails had died within 72 h. This result showed that *N. cumingii* was not able to tolerate heat stress.

### 4.2 Histology of the gills and kidney

Gills are the main respiratory organs of many aquatic animals, and they regulate osmotic pressure and excrete ammonia nitrogen (Qu et al., 2018). Andrew et al. (2011) found that in addition to salinity, environmental temperature can also affect the osmotic pressure balance and membrane permeability of the porgy fish (*Pagrus* sp.). In the current study, when the temperature decreased below 16°C, the osmotic pressure of the *N. cumingii* gill became imbalanced. Regaining balance required a lot of energy, so the number of mitochondrial cells and red blood cells increased. As the temperature dropped again, the blood vessels in the gills were constricted, and the uneven distribution of red blood cells was more evident. At 8°C, the blood vessels in the small gills gradually absorbed water and expanded, eventually swelling and flowing out of the red blood cells. This was similar to the results of Chen’s research on the gills of *Tegillarca granosa* (Chen et al., 2012). When the temperature dropped to 4 and then 0°C, the contraction of the gill lamella became more intense, and the number of red blood cells decreased. Similarly, the damage to gill cells in *Oreochromis niloticus* increased with decreased water temperature (Bing et al., 2015). *N. cumingii* cultured at 24°C exhibited edema in the gill lamella. At 28°C, the gill fragments were seriously deformed, and the distance between the gill capillary and the surrounding area was increased. These changes would have resulted in decreased gas exchange capacity and ultimately in damage to the snail. This result is similar to Cheng’s (2019) research about the structural changes of gill in scallop under high temperature.

The kidney is also an important excretory organ in the body of aquatic animals. Its function is to remove metabolic wastes from the body and regulate osmotic pressure (Shi et al., 2014). Li et al. (2013) reported that the kidney of the snail *Pomacea canaliculata* is composed of renal tubules and collecting ducts. In the current study, no obvious changes in the renal tubules and collecting ducts of the *N. cumingii* kidney were detected at different culture temperatures. However, the columnar cells were longest in the 16°C group. When the temperature gradually decreased or increased, the columnar cells became shorter and more numerous. This experiment was an acute test, and the temperature decreases or increases over a short time period likely accelerated the metabolic rate of the *N. cumingii* body.

### 4.3 T-AOC and activities of CAT and SOD

T-AOC is a comprehensive index used to assess the functional status of the body’s antioxidant system (De Oliveira et al., 2004; Tania et al., 2005; Guan et al., 2010). The antioxidant enzyme system mainly consists of SOD and CAT, which can decompose O_2_^−^ and H_2_O_2_. Xie (2016) studied the antioxidant capacity of the liver of *Pampus argenteus* under acute temperature stress and found that the T-AOC decreased, indicating that temperature stress caused a stress response in this species. In the current study, the T-AOC of the gills and kidneys of *N. cumingii* differed only slightly at temperatures between 4 and 24°C. In the gill, T-AOC first increased, then decreased, and then increased again as the culture temperature increased. This pattern differed from the results of most acute temperature stress experiments, and we speculate that the metabolic rate of the snail was accelerated quickly within a short period, and excessive reactive oxygen species (ROS) were produced, which intensified the oxidation reaction. As the temperature increased, T-AOC of the kidney first increased and then decreased, which is similar to the results reported by Feng et al. (2012) for the Chinese sturgeon *Acipenser sinensis*.

Water temperature can affect the metabolic rate of fish and the physiological activities of aquatic animals (Martinez et al., 2005). Li et al. (2008) set culture temperatures at 12, 21, 26, and 31°C and measured ROS content and SOD and CAT activities in the serum of Chinese sturgeons. All of these parameters increased with increasing temperature, which showed that the temperature increase likely promoted the production of ROS and the oxidation state of cell components, leading to reactions of related antioxidants and oxidase systems (Demple, 1999). In *N. cumingii*, the SOD and CAT activities of the gills first decreased and then increased with increasing temperature, which was similar to the activity of enzymes in the liver of *GIFT Nile tilapia* (Wang et al., 2012). However, as the temperature increased, the SOD activity of the *N. cumingii* kidney showed a W-shaped pattern, which was inconsistent with most results of the effect of temperature on the antioxidant defense system of aquatic animals. We speculate that this result was due to the acute nature of this experiment. When the experimental snails were exposed to external high or low temperature stress, the oxidative stress reaction caused the body’s metabolic rate to increase and excessive ROS to be produced within a short period of time. Too many ROS likely induced increased activities of SOD and CAT to eliminate the excessive O_2_^−^ and H_2_O_2_ in the body to maintain the balance of the snail’s antioxidant defense system. With increased temperature, the CAT activity in the *N. cumingii* kidney increased first, then decreased and then increased again, which was similar to the pattern of SOD and CAT activities in the liver of Gifford strains of *Nile tilapia juveniles* (Nurdiani et al., 2007).

### 4.4 Transcriptome analysis

In recent years, the survival of aquatic animals have been greatly threatened by the increase of global temperature (Brander, 2007), as heat stress is one of the most important environmental stress factors affecting aquatic animals (Zhou et al., 2018). Under stress conditions, the metabolism of aquatic animals depends greatly on environmental temperature. Studies have found that changes in the metaquatic animals abolic rate of the body are related to the increase in environmental temperature (Vergauwen et al., 2010; Qian et al., 2016; Jiang, 2013). Transcriptomics technology has provided a new way to study the effects of temperature on the immune response, growth, and development of aquatic organisms (Luo et al., 2015). In this study, it was found that the gill and kidney tissues of *N. cumingii* were significantly different at 16 °C and 28 °C through the study of feeding behavior, histology and immunoenzyme activity of *N. cumingii*. Therefore, sequencing was conducted at these two temperatures, the results showed that of the 4397 genes with differential expression in the gills of snails, 2339 were up-regulated and 2058 genes were down-regulated. Of the 4360 differentially expressed genes in the kidney of *N. cumingii*, 2300 were up-regulated and 2060 were down-regulated. We speculate that high temperatures may have activated cell metabolism and responded to the damage caused by a high temperature stress environment. Studies have shown that when the body is stressed, the structure and function of many enzymes and structural proteins change. For example, to protect the body from stress, it would stimulate the synthesis of heat shock proteins (Jiang, 2017; Hamdoun and Cherr, 2003).

The GO analysis of the gills and kidneys of *N. cumingii* showed that the number of down-regulated genes in both the high-temperature and the normal temperature group was significantly higher than the number of up-regulated genes. There was little difference in the number of up- or down-regulated genes in the cell composition and molecular function categories. This may be because when *N. cumingii* responds to high-temperature stress, its metabolism and energy consumption continue to increase, and the differentially expressed genes may respond to high-temperature stress through metabolic pathways. KEGG annotation statistics revealed that transcriptome differences were concentrated in six categories and 42 major metabolic pathways, and the largest difference was in the signal transduction process in the environmental information process. Studies have found that when the organism was under environmental stress, the function and structure of many enzymes and structural proteins will change. Meantime, the organism protects itself against adversity by up regulating the synthesis of heat shock proteins. Heat shock proteins played an important role in the synthesis, transport and glycosylation of proteins (Jayasundara et al., 2013; Ryckaert et al., 2010). In this study, The genes involved in the protein processing in the endoplasmic reticulum pathway have the highest proportion in the gills. The up-regulated genes mainly included Sec61, PDIs, Nef, HSP70 and Hsp90. The function of Sec61 is mainly to transfer the protein subunits or misfolded peptides to the cytoplasmic matrix for ubiquitination. PDIs play an important role in cell differentiation and the maintenance of function and cell activity (Liu, 2017). The up-regulating of HSP 70, HSP 90 and negative factor (Nef) indicates the feature of heat shock proteins expression under high temperature stress. In addition to heat stress proteins, we also found the immune-related gene TNF in the kidneys. Studies have shown that TNF signaling pathway is a multi effect pro-inflammatory cytokine. Meantime, TNF- α also participates in the regulation of cardiomyocyte apoptosis, and the signal transduction pathway mediated by it had a bidirectional effect, which can not only promote cardiomyocyte apoptosis, but also inhibit it (Petrus, 2011; ZHOU, 2010). Therefore, we suggested TNF plays an important role in the immune defense of *N. cumingii.*

The current study used high-throughput sequencing technology to sequence the *N. cumingii* transcriptome. Through the annotation of genes, we obtained a preliminary understanding of gene functions and a list of biological processes and metabolic pathways involved in the response of this snail to different culture temperatures. Our results provide valuable data for functional gene cloning, genomics, disease and stress resistance research, genetic breeding, and resource restoration.

## 5. Conclusion

In conclusion, *N. cumingii* was very sensitive to changes in temperature. High or low temperature could affect the tissue structure, immune enzyme activity and gene expression in gill and kidney tissues of *N. cumingii*. when the temperature was 8-16°C, *N. cumingii* had the best state with good food intake and active exercise status. Finally, we think the results obtained from this article can provide a good theoretical basis for the healthy culture and artificial reproduction of *N. cumingii* in the future.

## Acknowledgments

The authors wish to express thanks to the staffs of Key Laboratory of Mariculture & Stock Enhancement in North China’s Sea, Ministry of Agriculture, P.R.China for their help with the experiment. The authors are also grateful to the anonymous reviewers for the great elaboration of the manuscript through their critical reviewing and comments.

This study was supported by funds from National Natural Science Foundation of China (42076101).

## Notes

### Competing Interest Statement

The authors have declared no competing interest.

